# Generating an *in vitro* gut model with physiologically relevant biophysical mucus properties

**DOI:** 10.1101/2022.02.18.481062

**Authors:** Jacob McCright, Arnav Sinha, Katharina Maisel

## Abstract

**Introduction:** Gastrointestinal (GI) in vitro models have received lasting attention as an effective tool to model drug and nutrient absorption, study GI diseases, and design new drug delivery vehicles. A complete model of the GI epithelium should at a minimum include the two key functional components of the GI tract: mucus and the underlying epithelium. Mucus plays a key role in protecting and lubricating the GI tract, poses a barrier to orally administered therapies and pathogens, and serves as the microenvironment for the GI microbiome. These functions are reliant on the biophysical material properties of the mucus produced, including viscosity and pore size.

**Methods:** In this study, we generated *in vitro* models containing Caco-2 enterocyte-like cells and HT29-MTX goblet-like cells and determined the effects of coculture and mucus layer on epithelial permeability and biophysical properties of mucus using multiple particle tracking (MPT).

**Results:** We found that mucus height increased as the amount of HT29-MTX goblet-like cells increased. Additionally, we found that increasing the amount of HT29-MTX goblet-like cells within culture corresponded to an increase in mucus pore size and mucus microviscosity, measured using MPT. When compared to *ex vivo* mucus samples from mice and pigs, we found that a 90:10 ratio of Caco-2:HT29-MTX coculture displayed similar mucus pore size to porcine jejunum and that the mucus produced from 90:10 and 80:20 ratios of cells shared mechanical properties to porcine jejunum and ileum mucus.

**Conclusions:** GI coculture models are valuable tools in simulating the mucus barrier and can be utilized for a variety of applications including the study of GI diseases, food absorption, or therapeutic development.

**Biography:** Dr. Maisel joined the University of Maryland in January 2019 having done interdisciplinary training in nanotechnology, mucosal immunology, lymphatic immunology, and immunoengineering. She completed her PhD in Biomedical Engineering at the Johns Hopkins University in 2014 after which she was an NIH postdoctoral fellow at the University of Chicago in Molecular Engineering and Immunology. The Maisel Lab uses in vitro modeling, nanotechnology, and immunoengineering approaches to study and develop treatments for diseases at mucosal surfaces. They are interested in designing nanoparticles to take advantage of and study the interface between biological barriers, particularly the lymphatics, interstitial tissue, and mucosal surfaces, and nanoparticles. Dr. Maisel has won a number of awards, including NSF GRFP and NIH F32 fellowships as a trainee, the American Lung Association Dalsemer Award, LAM Foundation Career Development Award, an NSF CAREER Award, and an NIH NIGMS Maximizing Investigator Research Award. Her work has led to numerous high-impact publications, particularly in the field of drug delivery and mucosal immunoengineering, and several patents.

## Introduction

Mucus is the first line of defense to the outside world, covering our lungs, eyes, gastrointestinal tract, and urogenital tract. Mucus is a viscoelastic hydrogel-like network composed of water and mucins that effectively traps pathogens and foreign particulates, preventing them from harming or infecting the underlying epithelial cells. Mucins are high molecular weight glycoproteins, containing hydrophilic and hydrophobic domains that link and form a mesh with pores that can vary considerably between species of origin^1-4^ and fall within the range of 100-500 nm^5,6^. In addition to physically trapping larger materials, the hydrophobic domains and negatively charged glycosylation on mucins entrap positively charged and hydrophobic materials^7^. In the gastrointestinal tract (GI) mucus serves as a lubricant for food, and as a barrier against pathogens, digestive enzymes and acids, digested food particles, microbial by-products, food-associated toxins. Mucus also serves as the microenvironment for the commensal GI microbiome^8,9^. Understanding GI mucus structure is particularly important, as it is not only affected by disease, but can also modulate drug absorption – a key area of study to better design orally-delivered medications. However, it is difficult to access human GI tissues and thus access to human mucus samples has been extremely limited. In the last two decades, many *in vitro* models of the GI tract have been developed to both address questions about GI physiology, and to study therapeutic absorption and toxicity to improve drug and drug delivery designs.

The most traditional GI *in vitro* models use Caco-2 cells, an immortalized human colorectal adenocarcinoma cell line. These cells grow into confluent monolayers, differentiate into polarized enterocyte-like cells, and have been used extensively to screen for the permeability and toxicity of drugs^10-12^. Caco-2 Transwell® models have been shown to recapitulate the transport of insulin, calcitonin, and exenatide that is found in *ex vivo* models^13^. However, a main criticism of the use of these Caco-2 models^13^ is that, contrary to using the established 21-day culture period typically used to generate Caco-2 models, many reports cultured models for only 3-5 days prior to performing experiments, not allowing for differentiation into more representative GI epithelial cells^14^. Indeed, culturing conditions have been shown to massively affect how Caco-2 cells behave *in vitro*. Caco-2 cell models have been shown to increase in TEER and metabolic gene expression as passage number increases^15^. Also, studies on mannitol permeability across Caco-2 monolayers have demonstrated that permeability declines over culturing time, leveling off after 21 days, suggesting complete differentiation at this point^16^. Caco-2 cell layers have also been effective in demonstrating the toxicity of both new drug formulations, and environmental hazards, such as microplastics^17^. Collectively, Caco-2 Transwell® models have proven to be a valuable tool to model the absorptive qualities of the GI tract, but they lack the mucus layer that covers the GI epithelium which can modulate transport of materials toward it, thus affecting absorption.

Addition of a mucus layer is crucial to building physiologically relevant GI *in vitro* models^1,18^. In recent years, a new cell line has emerged, HT29-MTX, that form goblet-like cells which produce mucus. Co-culturing HT29-MTX cells with Caco-2 cells generates GI epithelial models with similar gene expression profiles to GI tissue, and results in a mucus layer on top of the epithelium^19^. Studies have examined how modifying the ratio of HT29-MTX cells to Caco-2 cells can change the properties of the epithelial layer, including the thickness of the mucus produced and the resistance across the GI model. Results of these studies have been relatively variable, generally indicating that the inclusion of HT29-MTX goblet cells results in a reduction of transepithelial resistance (TEER), and an increase in mucus height, measured using microscopy^20^. Mucus layers produced by these models have been found to be 10 – 50 μm in size^21,22^, and these coculture systems have largely been used to study how mucus affects drug and nutrient bioavailability^12^. Interestingly, cocultures with 10-20% HT29-MTX cells better recapitulate iron absorption trends, despite Caco-2 cells being the main contributor of iron transporters, suggesting that the inclusion of goblet-like cells modulates enterocyte physiology^23^.

While new GI *in vitro* models now usually include mucus-producing cells, characterization of the mucus in these model systems has been extremely limited. Typically, analysis of mucus from *in vitro* models consists of measuring electrical resistance, MUC2 (the primary mucin found in GI mucus) protein content, mucin gene expression in goblet cells, and imaging^24,25 26^. These methods are useful for confirming the presence of mucus; however, they provide relatively little information about the biophysical properties of the mucus produced by these cultures. The biophysical properties of mucus can affect its functions as a lubricant, mesh-like sieve, and homeostatic environment for the microbiome^27^. One key method to study the biophysical properties of mucus is rheology, but the limited volume of mucus produced by 1 cm^2^ or smaller Transwell® models, makes use of traditional rheometers difficult, as these require >100 μL of mucus^28^. Determining microrheology through multiple particle tracking (MPT) is a useful method to gain insight into the micron-level mechanics of mucus and can be performed using the low volumes produced by *in vitro* culture models^29,30^. MPT has already been used to determine pore sizes in vaginal, GI, and respiratory mucus^31-33^. In this study, we generated a coculture GI epithelial model using Caco-2 enterocyte-like cells and HT29-MTX goblet-like cells. We modified the ratio of the two cell types to simulate physiologically relevant cell numbers throughout the GI tract and determined its effect on the biophysical properties of the mucus within the model. We examined traditional hallmarks including epithelial morphology, TEER, and permeability, provided biophysical analysis of the mucus layer, and compared mucus properties to that of intestinal mucus from two common animal models: mice and pigs.

## Methods

### Cell Culture

Caco-2 (p44-48) and HT29-MTX cells (p32-40) were cultured in 75-cm^2^ plastic flasks (Nunc EasyFlask Thermo Scientific, Nuclon Delta Surface). Culture medium consisted of Dulbecco’s Modified Eagle’s Medium with 10% Fetal Bovine Serum (Gibco) for both Caco-2 and HT29-MTX lines. Coculture of Caco-2 and HT29-MTX was performed at the following ratios of Caco-2:HT29-MTX: 1:0, 9:1, 4:1, and 0:1. Cells were seeded at a total density of 375,000 cells/cm^2^ in 1.0 μm pore size cell culture inserts (PET, Falcon) in a 12 Well Tissue Culture Plate (Falcon). Cocultures were maintained for 21 days to allow Caco-2 cells to differentiate and develop enterocyte-like features. All cultures were kept at 37°C in 5% CO_2_, 95% air atmosphere and regularly checked every other day to change medium. Mucus was washed from cultures using phosphate buffered saline (PBS) weekly. Trans epithelial electrical resistance (TEER) was measured on day 7, 14, and 21 using a millicell ERS-2 voltohmmeter (Millipore Sigma). Viability was assessed through staining with propidium iodide (PI; 81845, Sigma). PI was administered to live cultures after 21 days of culture on 24 well plates (Nunc Thermo Scientific) at a concentration of 30 μM in PBS for 15 minutes in the dark. Cells were stained with propidium iodide and imaged using fluorescence microscopy to identify non-viable cells.

### Immunofluorescence Staining

Coculture Transwell© inserts were fixed with 4.0% cold Paraformaldehyde (PFA) in PBS for 15 min. Samples were then washed 3 times with PBS and kept at 4°C. Membrane inserts were cut out and placed cells-up on microscope slides. Samples were permeabilized using 0.1% triton X-100 for 15 minutes. Samples were blocked with 2% FBS in PBS for 30 minutes and then washed with PBS times. Primary antibody (anti-EpCAM, anti-MUC2 abcam) was diluted 1:200 in blocking solution and applied to samples for 2 h at room temperature. Primary antibody was then removed, and the sample was washed twice with PBS containing 0.1% Tween 20 and 3 times with PBS. Secondary antibody and phalloidin-AF555 (Invitrogen) were applied for 2 h at room temperature (Invitrogen). Cover slips were mounted on the samples with Vectashield Antifade Mounting Medium with DAPI and sealed with nail polish prior to imaging. Fluorescence imaging was performed using an Olympus FV300 Laser Scanning Confocal Microscope. Mucus height was determined by measuring the distance the fluorescently labeled mucus layer spanned using ImageJ (NIH). Briefly, XZ confocal projections stacks were averaged over the Y axis. Mucus ROI was traced and average height was recorded for each confocal image. 6 images were taken from n = 4 samples.

### Coculture Permeability

Cocultures were generated on 1.0 μm transwells and cultured for three weeks as described above. On day 21 10 μg/mL fluorescent 150 kDa Dextran, 4 kDa Dextran, or AF647-labeled albumin (Invitrogen) was added to the top well and transport to the bottom well was measured via fluorescence (Tecan Spark microplate reader) hourly for up to 6 h. Fluorescence intensity was quantified using a plate reader and tracer transport was calculated using a standard curve. Effective permeability was estimated using the equation: 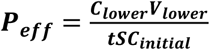, where C = concentration, V_lower_ = volume of the bottom compartment, S = surface area, and T = time.

### Microrheology

Non-mucoadhesive nanoparticles were generated using previously published methods^33-35^. Briefly, commercially available fluorescent 100 nm, carboxyl-modified polystyrene nanoparticles (Invitrogen) were coated with 5 kDa polyethylene glycol (PEG, Creative PEGworks) using EDC/NHS chemistry. PEG coating was confirmed using dynamic light scattering (DLS, Brookhaven Scientific). Mucus was collected from the surface of the transwells and 0.5 μL of nanoparticles at 1E-5 w/w%, or 1 μg/mL. All tracking was performed on day 21 of coculturing using an Axio Observer fluorescence microscope. Images were recorded at a frame rate of 30 frames per second. Centroid tracking was performed using automated MatLab software package based on a previously developed method ^36^. Briefly, the 2D localization of fluorescent nanoparticles was determined based on an intensity and eccentricity threshold. Trajecotries were determined by connecting particle centers between serial images. The Mean Squared Displacement (MSD) can be calculated as: <Δr2(T)> = <[x(t + τ) – x(t)]2 + [y(t + T) + y(t)]2>, where T is the time lag between frames and angle brackets denote the average over the time points of interest. The MSD was then used along with the Stokes-Einstein equation to calculate the viscosity: MSD=4Dt; D=k_B T/6πηr where: D = Diffusivity Constant, k_B_ = Boltzmann’s, Constant, r = Radius of nanoparticle, T = temperature, t = time, and η = Viscosity. Using these equations, we can determine the viscosity of the mucus. Further viscoelastic properties of the mucus was determined using the generalized Stokes-Einstein equation which relates the viscoelastic spectrum [G(s)] to the Laplace transform of <Δr2(T)>, <Δr2(s)>, with the equation G(s) = 2kBT/[πas<Δr2(s)>], where kBT is thermal energy, a is particle radius, and s is the complex Laplace frequency^29^. Making the substitution *s* with *i*ω where i is a complex number and ω is frequency, the complex modulus can then be calculated as G*(ω) = G’(ω) + G”(iω). The viscous and elastic moduli, G’’ and G’ can be calculated using the expression for complex microviscosity, η*(ω) = G*(ω)/ω. The pore size of the mucus hydrogel (ξ) can be estimated based on measured G’ as ξ ≈ (kBT/G′)1/3 ^37,38^.

For animal studies, female C57Bl/6J mice 8-10 weeks old were purchased from Jackson Labs and allowed to acclimate for up to 14 days. Porcine intestine samples were obtained from Animal Biotech Industries (ABI, Doylestown PA). Animal tissues were harvested and 5 μL of 1.0 μg/mL nanoparticles were added to the mucus covering the epithelial surface, without freezing or washing tissue to preserve mucus integrity. Microrheology was performed as described above. All animal studies were approved by the UMD IACUC.

### Statistics

Group analysis was performed using a 2-way ANOVA, followed by Tukey’s post-test. Unpaired Student’s t-test was used to examine differences between only two groups. A value of *p* < 0.05 was considered significant (GraphPad). All data is presented as mean ± standard error of the mean (SEM). All experiments were performed at least twice with 2-3 technical replicates per condition. Only statistically significant or trending (p ≤ 0.1) data is indicated in the Figures.

## Results

We sought to generate a model GI epithelium with physiologically relevant mucus by coculturing HT29-MTX mucus producing goblet-like cells with Caco-2 enterocyte-like cells. Caco-2 cells and HT29-MTX cells were seeded at four different ratios: 1:0, 9:1, 4:1, and 0:1 Caco-2:HT29-MTX (CH10:0, CH9:1, CH8:2, CH0:10, respectively), representative of the jejunum and ileum of the small intestine ^39^. Cells were cultured for 21 days on a 1.0 um Transwell© insert at confluence to allow differentiation of Caco-2 cells into enterocyte-like phenotype. As expected, Caco-2 cells did not secrete mucus, and no secreted mucus layer was observed (**Figure 1A**) ^40^. We found that f-actin is localized to cell membranes, particularly along the apical surface of the epithelial layer, and is most evident in cultures containing Caco-2 cells (**Figure 1A, Sup 1**). As HT29-MTX cells are added, the signal and continuity of the f-actin is reduced, but monolayer integrity is maintained, as no obvious gaps between cells can be observed. We found that the MUC2 secreted mucus layer increases in height as the amount of HT29-MTX cells in the culture is increased (**Figure 1A-B**). The mucus layer height is tallest on the cultures containing only HT29-MTX cells: 175 ± 37 μm, and smaller in the cocultures: 48 ± 13 μm for CH9:1, and 94 ± 10 μm for CH8:2 (**Figure 1B**). It is difficult to differentiate Caco-2 and HT29-MTX cells, since both express mucin and other typical GI epithelial genes/proteins, and thus it is possible that the original 9:1 and 8:2 ratios have changed throughout our culture even though cells are seeded at confluence. However, the changes we see in mucus accumulation, and f-actin signal suggest that even if the original ratios are not maintained after 21 days, the HT29-MTX cell population is different between the two conditions. Coculture viability was close to 100% for all conditions (**Figure 1C**).

**Figure 1:**
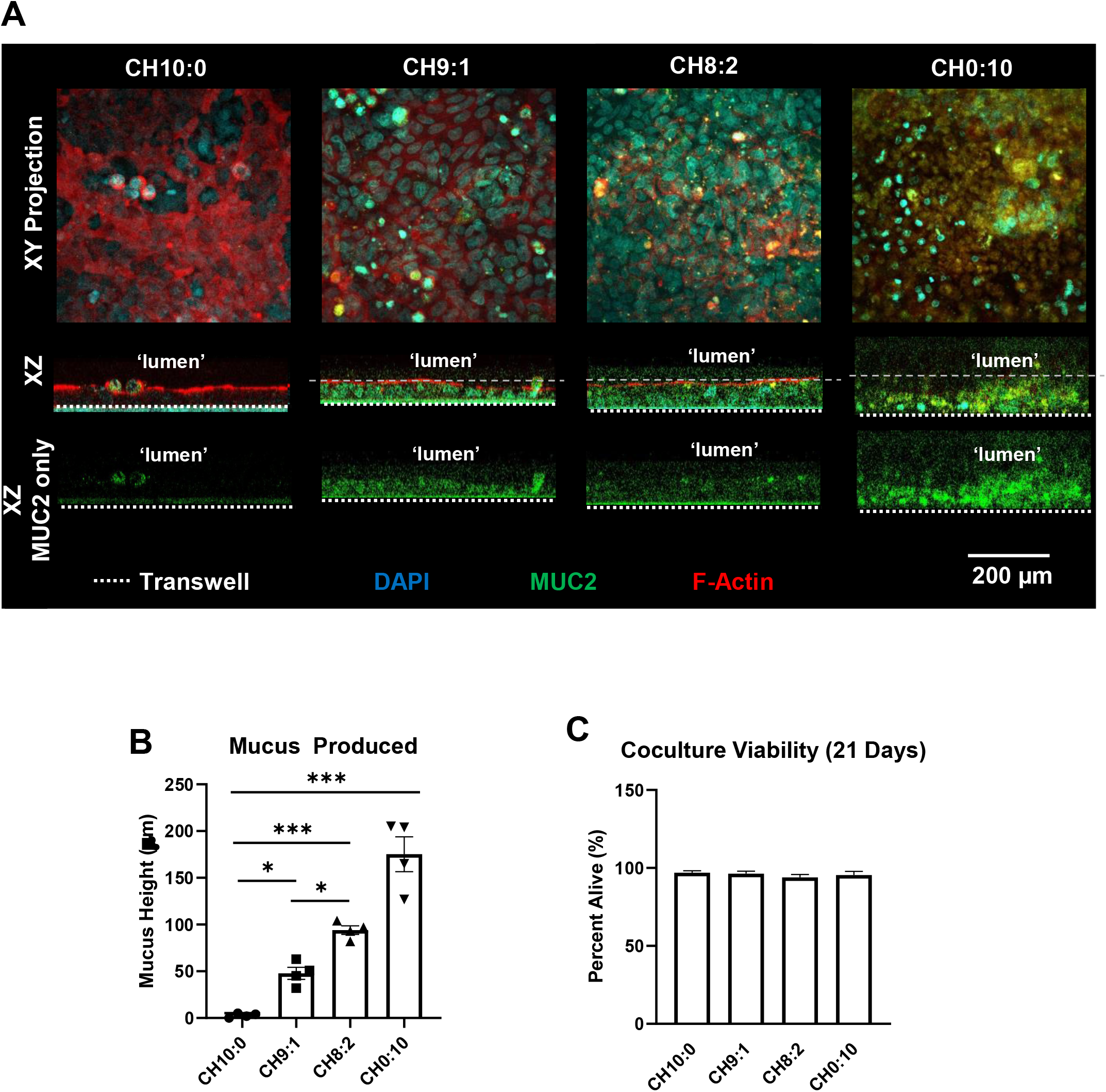
Morphology and Physiology of GI Co-Culture Models. (A) Immunofluorescence images of different co-culture conditions visualizing cytoskeletal F-actin (Phalloidin, red), mucin (MUC2, green) and nuclei (DAPI, blue). Grey dashed lines indicate representative location from which mucus height was measured and white dashed lines indicate location of the transwell membrane. B) Quantification of mucus head produced. C) Viability of coculture models at maturity (21 days). n = 4 (*P < 0.05, *** P < 0.001) measured using 1-way Anova. Scale bar indicates 200 μm.

After characterizing the morphology of the different GI cocultures, we sought to assess the effect of a mucus layer and HT29-MTX cells on model GI epithelial permeability. We found that TEER increased from 330 ± 30 to 440 ± 110 mV to 510 ± 60 mV for CH10:0 and from 180 ± 20 to 260 ± 30 mV to 230 ± 40 mV for CH0:10 (**Figure 2A**). Both mixed cultures also increased over time, with the CH9:1 culture increasing from 370±10 to 530±30 mV to 570 ± 50 mV and the CH8:2 culture increasing from 420 ± 20 to 550 ± 20 mV to 610 ± 40 mV (**Figure 2A**). We observed an increase in TEER with the addition of HT29-MTX cells to the culture across all time points. We also assessed solute permeability of the coculture model using fluorescent bio-inert 150 kDa dextran, 4 kDa dextran, and albumin, a model protein. Using the 4 kDa dextran tracer, commonly used in studying epithelial permeability, we observed that the HT29-MTX monoculture was the most permeable with 9.1 ± 0.8 % transport. This was significantly higher than the CH9:1 and CH8:2 cocultures where the 4 kDa transport efficiency was 3.5 ±0.2% and 4.0 ± 0.6% respectively. The Caco-2 monoculture was the least permeable to 4 kDa dextran with a transport efficiency of 1.5 ± 0.1% (**Figure 2B)**. The other coculture conditions, CH10:0, CH9:1, and CH8:2, displayed similar transport efficiency, ranging from 2.5 ± 0.1% - 2.5 ± 0.2% after 6 hours. Measuring the trans-epithelial transport of the 150 kDa dextran, we found that the uniform HT29-MTX culture was the most permeable to this larger molecule, with 3.0 ± 0.6% transport efficiency (**Supp 2**). Albumin transport was similar for all Caco-2 containing samples, though there was a trend toward lower permeability of the CH9:1, and the HT29-MTX culture was most permeable with 5.7±3.7% of the albumin transported after 6 h (**Figure 2C**). Our data demonstrate that the effective permeability for 4kDa dextran was the highest for the HT29-MTX monoculture (Figure 2D, Supp 2), and the other cell compositions displayed similar effective permeabilities for the different tracers (**Figure 2D-E, Supp 2**).

**Figure 2:**
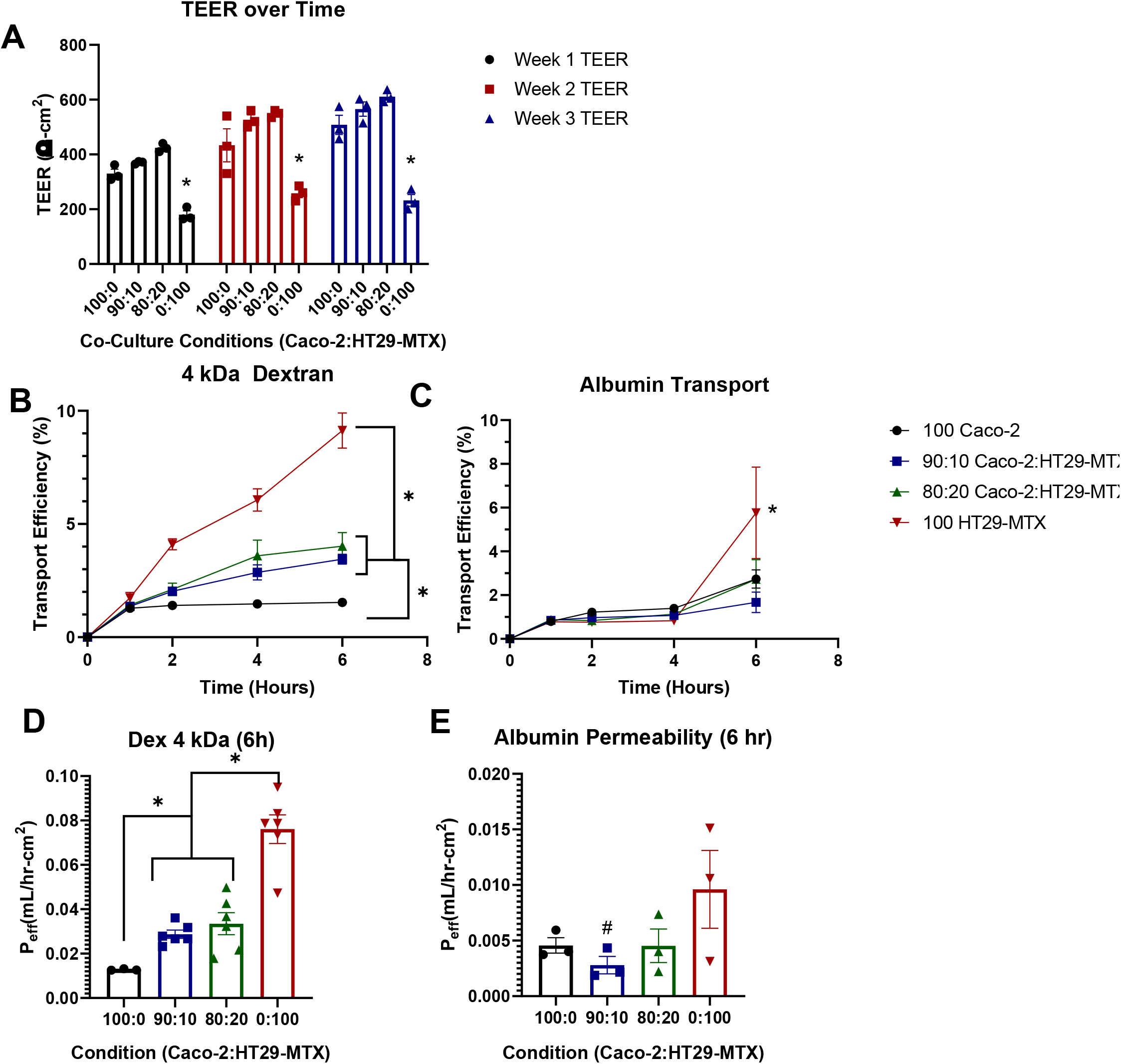
Permeability of GI Co-Cultures. A) Trans-Epithelial Electrical Resistance (TEER) of co-cultures over time. B) 4 kDa dextran and C) albumin transport efficiency across the transwell coculture over time after 3 weeks. Calculated permeability of cocultures at 6h to D) 4 kDa Dextran and E) albumin after 3 weeks. n = 3 – 6 (*p < 0.05, #p < 0.1) measured using 2-way Anova.

Next, we characterized the rheological properties of the mucus layer in our coculture model systems using MPT. As the Caco-2 cell lines do not secrete a mucus layer, we did not measure rheological properties of the CH10:0 coculture model. We found that MSD and diffusivity increased as the ratio of HT29-MTX cells within the culture increased. PEGylated non-mucoadhesive nanoparticles (MPP) displayed a diffusivity of 0.11 ± 0.2 μm/s^2^ in CH9:1 mucus, and 0.12 ± 0.01 μm/s^2^ in CH8:2 mucus. MPPs within mucus samples from goblet-cell monocultures (CH0:10) had a diffusivity of 0.13 ± 0.01 μm/s^2^ (**Figure 3A**), indicating that all cultures had similar diffusivities. We found that the viscosity of the CH9:1 mucus was 0.03 ± 0.01 Pa-s (**Figure 3B**). The mucus of the CH8:2 mucus was ∼25% more viscous than CH9:1 mucus, at 0.04 ± 0.01 Pa-s (**Figure 3B**), while CH0:10 mucus was the most viscous at 0.045 ± 0.01 Pa-s (P<0.2), though none of these differences were statistically significant (**Figure 3B**). We then extrapolated pore size of the mucus layers and found that there was a relatively large variability. We found that the average pore size was 440 ± 15 nm^2^ in the CH9:1 mucus, 610±30 nm^2^ in the CH8:2 mucus, and 620 ± 70 nm^2^in the CH0:10 mucus, with CH9:1 mucus having a significantly lower pore size than both CH8:2 and CH0:10 (**Figure 3C**). However, pore sizes ranged from 10 to 1200 nm^2^ for CH9:1 and 10 to 2800 nm^2^ for CH8:2, highlighting the heterogeneity of the produced mucus layer (**Figure 3C**). Finally, we determined the elastic (storage) modulus (G’) and plastic (loss) modulus (G’’). G’ at τ=0.5s was similar at 0.15 ± 0.06 Pa and 0.20 ± 0.12 Pa (ns) for CH9:1 vs. CH8:2, respectively (Figure 3D). The monoculture of HT29-MTX goblet-like cells had the highest elastic modulus of 0.48 ± 0.19 Pa, though due to the high variability this was only a trend when compared to CH9:1 (**Figure 3D**). The plastic modulus and ratio between G’/G” was highest for HT29-MTX monocultures (**Figure 3E-F**). Interestingly, when measuring G’’/G’ we saw that the mucus produced by the cocultures behaved elastically (G’’/G’ < 1), whereas the monoculture of HT29-MTX cells was slightly plastic (G’’/G’ ≥ 1) (**Figure 3F**).

**Figure 3:**
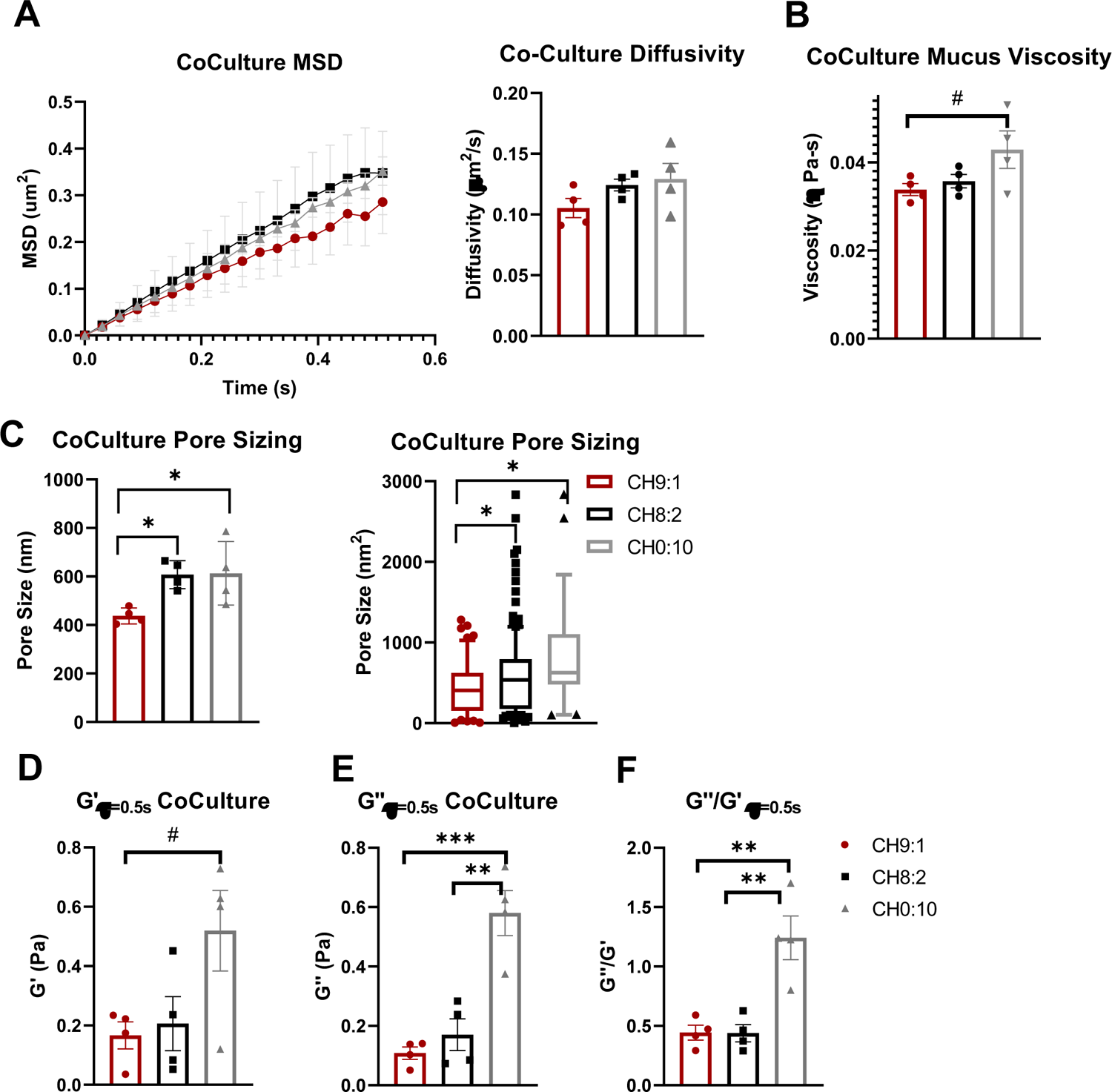
Material Properties of Mucus Produced by Coculture Models after 3 weeks. A) Diffusivity of 100 nm PEGylated nanoparticle in culture-produced mucus samples. B) Microvisocisty, C) pore size D) elastic modulus (G’), E) plastic modulus (G’’), and F) G’’/G’ of mucus produced by cocultures after 3 weeks. n = 4-6 (#P < 0.1; *P < 0.05;) measured using 2-way ANOVA for MSD and 1-way ANOVA otherwise.

To determine if the properties of the mucus produced in our *in vitro* model was representative of *in vivo* mucus, we collected GI mucus from mice and pigs, two animal models commonly used for studying the GI tract. We compared the mucus properties in both the jejunum and ileum of the small intestines. We found that there was a trend toward higher MPP diffusivity within the jejunum when compared to the ileum for both porcine and murine mucus samples (**Figure 4A**). MPP diffusivity was 0.40 ± 0.1 to 0.20 ± 0.1 μm/s^2^ for the porcine samples and 0.30 ± 0.1 to 0.19 ± 0.1 μm/s^2^ for the murine mucus samples. We also found that porcine jejunal mucus had a viscosity of 0.034 ± 0.01 Pa-s, less viscous than porcine ileal mucus, which had a viscosity of 0.087 ± 0.01 Pa-s (**Figure 4B**). The porcine jejunal mucus displayed similar microviscosity measurements to both coculture models 0.036 ± 0.002 Pa-s and 0.034 ± 0.001 Pa-s for CH9:1 and CH8:2, respectively. Murine mucus viscosity was higher in the jejunum compared to porcine jejunum, measuring 0.07±0.02 Pa-s in the murine jejunum, and was similar to porcine ileal mucus, measuring 0.10 ± 0.03 Pa-s in the murine ileum (**Figure 4B, Table 1**). We found that pore size was consistently below 500 nm^2^ for all porcine and murine small intestinal mucus samples (**Figure 4C**). Porcine jejunal mucus had a mucus pore measuring 390 ± 50 nm^2^, similar to the mucus produced by the CH9:1 coculture model (440 ± 15 nm^2^) (**Figure 4C**). Calculating the material properties of the animal mucus, we found that both were elastic in nature, evidenced by the ratio of G’’/G’ < 1 (**Figure 4D-F**), similar to our coculture in vitro models. Collectively, these results indicate that there are similarities in material properties between the mucus produced by the *in vitro* models and mucus from *in vivo* animals (**Table 1**). Additionally, we identified that the ratio of 90:10 Caco-2 to HT29-MTX cells recapitulated the viscosity, pore size, G’ and G” of porcine jejunal mucus (**Table 1**).

**Table 1:** Summary of material properties of different co-culture systems compared to pig and mouse mucus. Values given as mean ± SEM (n=3-6) measured using 1-way Anova. Symbol (& for jejunum or @ for ileum, mouse – blue, pig – red) indicates that the measured parameter (viscosity, etc) for the coculture model is not statistically different or trending toward a difference compared to the respective animal mucus (p > 0.1).

|  |  | Viscosity<br>(Pa-s) | Pore Size<br>(nm<sup>2</sup>) | G'<br>(Pa) | G''<br>(Pa) | G'/G'' |
| --- | --- | --- | --- | --- | --- | --- |
| **Caco-2:**<br>**HT29-**<br>**MTX** | **9:1** | 0.034 ±<br>0.001 & | 440 ± 50 & | 0.16 ± 0.05<br>&, @, &<br>@ | 0.11 ± 0.02<br>& | 0.44 ± 0.06 |
|  | **8:2** | 0.036 ±<br>0.002 & | 600 ± 40 | 0.21 ± 0.09<br>&, @, &<br>@ | 0.17 ± 0.05<br>& | 0.44 ± 0.07 |
|  | **0:10** | 0.043 ±<br>0.004 | 830 ± 120 | 0.52 ± 0.13 | 0.58 ± 0.07 | 1.24 ± 0.18 |
| **Mouse** | **Jejunum**<br>**(&)** | 0.065 ±<br>0.009 | 220 ± 30 | 0.25 ± 0.05 | 0.21 ± 0.02 | 0.86 ± 0.09 |
|  | **Ileum**<br>**(@)** | 0.100 ±<br>0.032 | 180 ± 10 | 0.22 ± 0.01 | 0.18 ± 0.02 | 0.8 ± 0.02 |
| **Pig** | **Jejunum**<br>**(&)** | 0.034 ±<br>0.009 | 390 ± 30 | 0.3 ± 0.04 | 0.25 ± 0.04 | 0.83 ± 0.08 |
|  | **Ileum**<br>**(@)** | 0.087 ±<br>0.008 | 300 ± 50 | 0.26 ± 0.02 | 0.2 ± 0.01 | 0.8 ± 0.1 |

**Figure 4:**
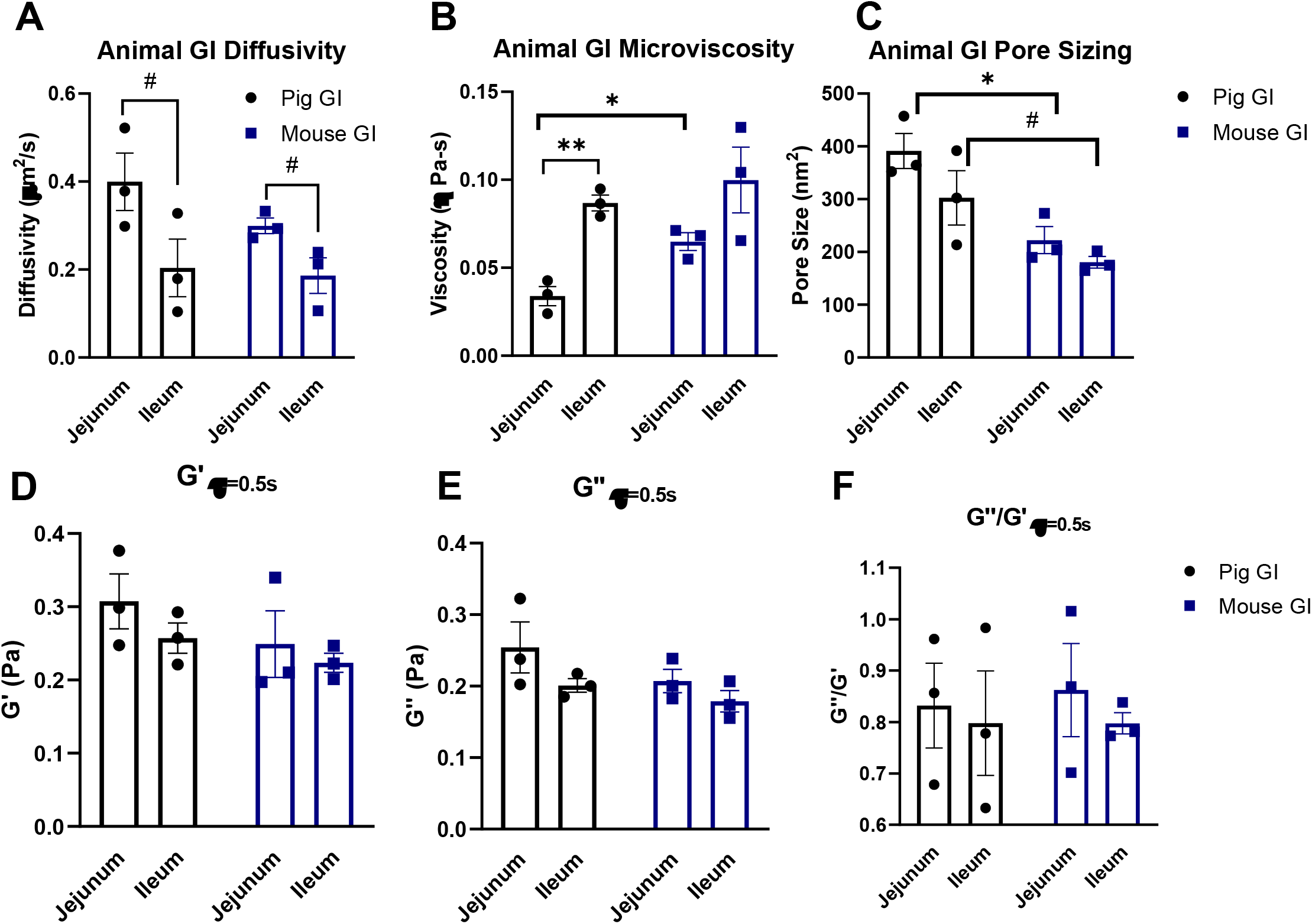
Material Properties of Mucus from In-Vivo Animal Models. A) Diffusivity of 100 nm PEGylated nanoparticle in collected mucus samples. B) Calculated microvisocisty of mucus samples. C) Pore size of mucus samples D) elastic modulus (G’), E) plastic modulus (G’’), and F) G’’/G’ of mucus. n = 3 (#: p = 0.054; *p < 0.05) measured using Student’s t-tests.

## Discussion

In this work we established an *in vitro* model of the mucus-producing GI epithelium and characterized the permeability of the model as well as the material properties of the mucus produced by the model. We found that increasing the number of HT29-MTX cells within our culture model resulted in more mucus production, as indicated by an increase in mucus height. The addition of HT29-MTX cells, and thus a mucus layer, resulted in an increase in TEER, but did not significantly affect the model’s permeability of the albumin and 4 or 150 kDa dextrans compared to Caco2 cells alone. Our microrheological analysis revealed that the mucus became more viscous and contained larger pores as the amount of HT29-MTX cells was increased. Interestingly, we show that the mucus produced by the coculture was distinct from the mucus produced by the HT29-MTX cells alone, as the mucus produced by the cocultures was more viscoelastic than that produced by the monoculture of HT29-MTX cells. The mucus produced by the coculture models was also less viscous than that found in both the porcine and murine GI mucus. In summary, we generated and characterized an *in vitro* model for the GI epithelium that can inform oral drug delivery strategies while providing some of the first analyses of the mucus layer produced by an *in vitro* GI model.

Studies have found that the addition of a mucus layer via coculture of HT29-MTX and Caco-2 cells does not modulate permeability of the *in vitro* model system compared to Caco-2 cells alone^42-44^. Our study confirms these findings: We found that changing the ratio of cells within our culture did not change the permeability of 4 or 150 kDa dextran and albumin. Others have also reported that the addition of HT29-MTX cells does not change the permeability of chemical, macromolecule, or protein tracers in a significant way ^42,44^. However, permeability is increased in monocultures of HT29-MTX cells, likely due to their lack of forming adequate tight junctions. Similarly, literature suggests that addition of HT29-MTX cells modifies formation of tight junctions between the enterocyte-like cells, which has been observed in numerous other studies using coculture ratios ranging from 70:30 – 90:10 Caco-2 to HT29 cells, and occurs because the goblet-like cells do not adequately form tight junctions themselves^20,45,46^. Despite this, other groups found that changing the ratios of the cell types did not alter the TEER values measured^46,47^. This is contrary to what we have observed, where the introduction of HT29-MTX cells seems to have increased the TEER of our models. This inconsistency may be caused by the thicker mucus layer in our system, which may alter the TEER readings. Despite the differing trends in TEER as a result of the addition of HT29-MTX cells, TEER values for cocultures range from 300-900 Ω-cm^2^, which is consistent with our models. Another consideration for models utilizing differing ratios of Caco-2 and HT29-MTX cells is that due to the challenge of definitively identifying them through imaging, it can be difficult to establish that the original ratio of cells are maintained due to different proliferation rates. Overall, our system recapitulates the permeability and mucus production of previously developed *in vitro* model systems.

Mucus height changes throughout the different parts of the GI tract, which is unsurprising since the jejunum, ileum, and colon have distinct functions – food digestion/absorption in the small intestine and water/ion absorption in the colon. The duodenum has the thickest mucus layer, measuring up to 500 microns^27^, likely to prevent the incoming gastric juices and digestive enzymes from reaching the epithelial surface too quickly. In the jejunum, mucus thickness is only 150-300 microns, likely to aid in food absorption, while the mucus layer in the ileum can measure up to 500 microns in thickness ^21^. These values are much greater than those recreated in classic *in vitro* models using 70:30 – 90:10 ratio of Caco-2 to HT29-MTX coculture models of the GI tract, where mucus thickness mostly resembles that of the firmly adherent mucus layer, ranging from 15 – 30 microns in the small intestine^3,22,24,48^. Navabi et al completed a thorough study where they examined a combination of Caco-2 and HT29-MTX ratios and strains, and the effect of mechanical stimulus on mucus production. The mucus layer produced by their models was consistently thinner than that of our model^48^. Our model system had mucus of 40 – 80 microns, suggesting that our system includes a thicker mucus layer than previously reported systems, which may make it more physiologically representative. Further optimization to enhance the height of the mucus layer *in vitro* may be needed to build truly representative *in vitro* model systems.

In addition to mucus height, the material properties of the mucus change along the different parts of the intestine. Studies in mice have suggested that mouse colonic mucus has an average pore size close to 200 nm, as 200 nm, non-mucoadhesive nanoparticles had reduced diffusivity in the mucus layer^33^. Mouse small intestinal mucus, on the other hand, had a larger pore size ranging between 200 – 500 nm, as 200 nm nanoparticles freely diffused with the mucus gel, while 500 nm nanoparticles did not^32^. Studies using reconstituted porcine-derived mucin have estimated that the pore size of small intestinal mucus lies between 100 – 200 nm in size^49^. This has been corroborated by Yildiz et al. who observed a 20-fold decrease in particle diffusivity when nanoparticle diameter was increased from 100 nm to 500 nm, suggesting that the pore sizes within porcine mucus lie between 100 – 500 nm^50^. A confounding factor in many of these studies is that the reliance on mucus hydrogels generated from purified and heavily processed gastric mucins, instead of small and large intestinal mucins, which can lead to gels that do not necessarily recapitulate the mucin seen *in vivo*^51^. Indeed, electron micrographs of *ex vivo* mucus highlight the heterogeneity of mucus pore sizes *in vivo* and provide a visual estimation of pore size closer to 500 nm diameter^52^. One group demonstrated that microparticles of 0.5 – 2 μm^3^ were able to diffuse through murine ileal mucus samples, suggesting that mucus pore sizes could be as large as 2 μm^2 53^. Our data shows that coculture in vitro models produce mucus with pore sizes more in line with that of estimated electron microscopy images of human mucus, a range of 50 - 300 nm in diameter^43^. Mucus pore size measurements have continued to result in seemingly conflicting data, which can largely be contributed to the myriad factors that can influence mucus permeability, including local concentrations of mucins and other macromolecules, fluid flow, presence of food materials, and diet. Collectively, these studies have suggested that pore sizes in mucus are extremely heterogeneous, which is recapitulated in our coculture models.

Mucus viscosity is key in maintaining barrier properties and is typically a function of pore size, with a tighter mesh network resulting in more viscous mucus. We found that the mucus from our cocultures had a viscosity of 0.03 - 0.04 Pa-s. This viscosity was below the reported ranges of 0.06 - 0.10 Pa-s in pig mucus, as measured via macrorheology^54^. However, differences in macro- and microrheology are common in other tissues as well. A similar study examining mucus barrier properties within different regions of the mucus layer corroborated our measurements of mucus microviscosity. Using MPT, they found that *ex vivo*, unmodified, jejunal porcine mucus had a microviscosity in the range of 0.01-0.05 Pa-s, and remained relatively similar at different depths of the mucus layer^28,32,55^. Despite these studies, how viscosity and pore sizes vary along the small intestinal tract is still largely understudied. Comparisons between small intestine mucus and colonic mucus suggest that viscosity of mucus increases as the GI tract progresses. Indeed, a study by Swidsinski et al. indirectly measured this gradient in mucus through analysis of the microbiome components within different regions of the GI tract^56^. Using agarose gel of different viscosities, they demonstrated that bacterial shape correlated with an environmental viscosity best suited for survival and mobility. Through this, they hypothesized that the shape of bacteria in specific regions of the GI tract can predict the material properties of that region. Using this strategy, they further hypothesized that mucus viscosity increases from proximal to distal regions of the intestines. This trend is corroborated by our findings where our *ex vivo* microrheological measurements of porcine and mouse mucus indicate that viscosity increases from the jejunum to the ileum.

With inflammatory diseases at the forefront of *in vitro* modeling using Caco-2 and HT29-MTX cultures, many groups are also experimenting with adding immune cells into GI models. This is typically achieved through the addition of Raji B cells, microfold (M) cells, as well as THP-1-derived macrophages^57,58^. Ude et al. incorporated Raji B cells within a Caco-2-based model of the GI tract to generate models sensitive to copper oxide pollutants^59^. Kerneis et al. added lymphocytes from Peyer’s patches to a Caco-2 GI model. Within the model, these lymphocytes behaved like M cells, transporting inert nanoparticles and bacteria^60^. Beterams et al. recently developed a triple coculture model consisting of THP-1-derived macrophages, T84 epithelial cells, and LS-174T goblet cells^61^. They found that the inclusion of immune cells resulted in an increase in mucus height from 60 to 80 μm, compared to a coculture with no macrophages, suggesting that inclusion of immune cells is crucial to create a more physiologically representative mucus height. The group further introduced model microbiota which induced pro-inflammatory cytokine production. While these studies highlight the promise of modeling both microbiota and disease interactions *in vitro*, a key next step for these models will be to include the microbiota within a mucus gel in these models. This is incredibly challenging as most microbiota require anaerobic conditions, while epithelial cultures require oxygen. However, generating mucus models with physiologically relevant material properties could facilitate the incorporation of microbiota and allow them to behave as they would *in vivo*.

In this study we demonstrated how coculture of enterocyte and goblet-like cells within a GI model resulted in physiologically relevant recapitulation of intestinal mucus properties. We found that increasing the number of goblet cells resulted in a more resistant monolayer, while also producing mucus that was more viscous with larger pores. Being able to accurately recapitulate both epithelium and mucus microenvironment is key for engineering oral therapeutics which must cross this barrier, as well as studying microbiome interactions with its physiologically relevant microenvironment. Future GI *in vitro* models should consider the material properties of the mucus produced, as features like pore size dictate microbiome health as well as drug delivery. Models can include additions such as immune cells and microbes to provide more physiologically relevant model systems.

## Acknowledgements

We would like to thank the UMD Bioworkshop for core facility usage and Michele Kaluzienski for assisting with manuscript preparation. This work was funded by the University of Maryland New Directions Fund and Faculty-Student Research Award (KM) and the AHA Predoctoral Research Fellowship (JM).

---

**Supplemental Figure 1:**
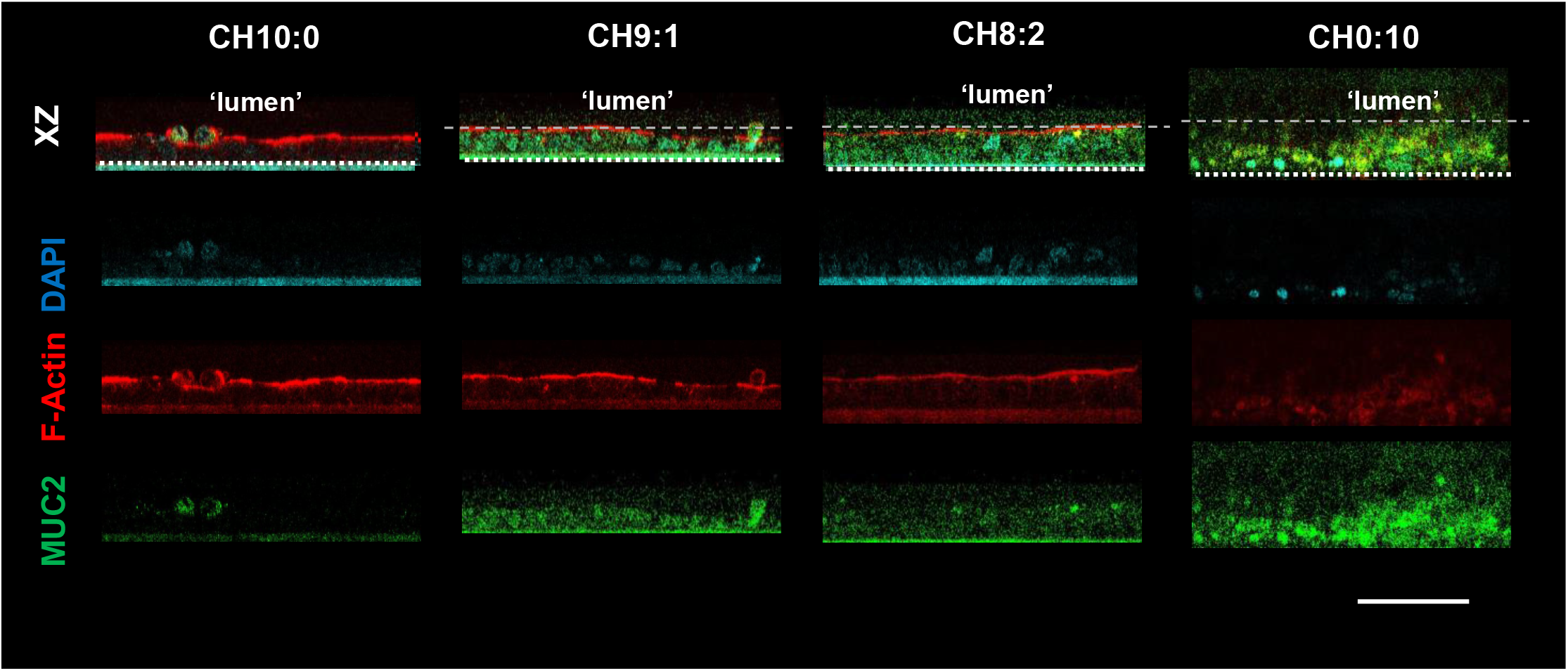
(A) Immunofluorescence images of different co-culture conditions visualizing cytoskeletal F-actin (Phalloidin, red), mucin (MUC2, green) and nuclei (DAPI, blue) Scale bar 200 μm.

**Supplemental Figure 2.**
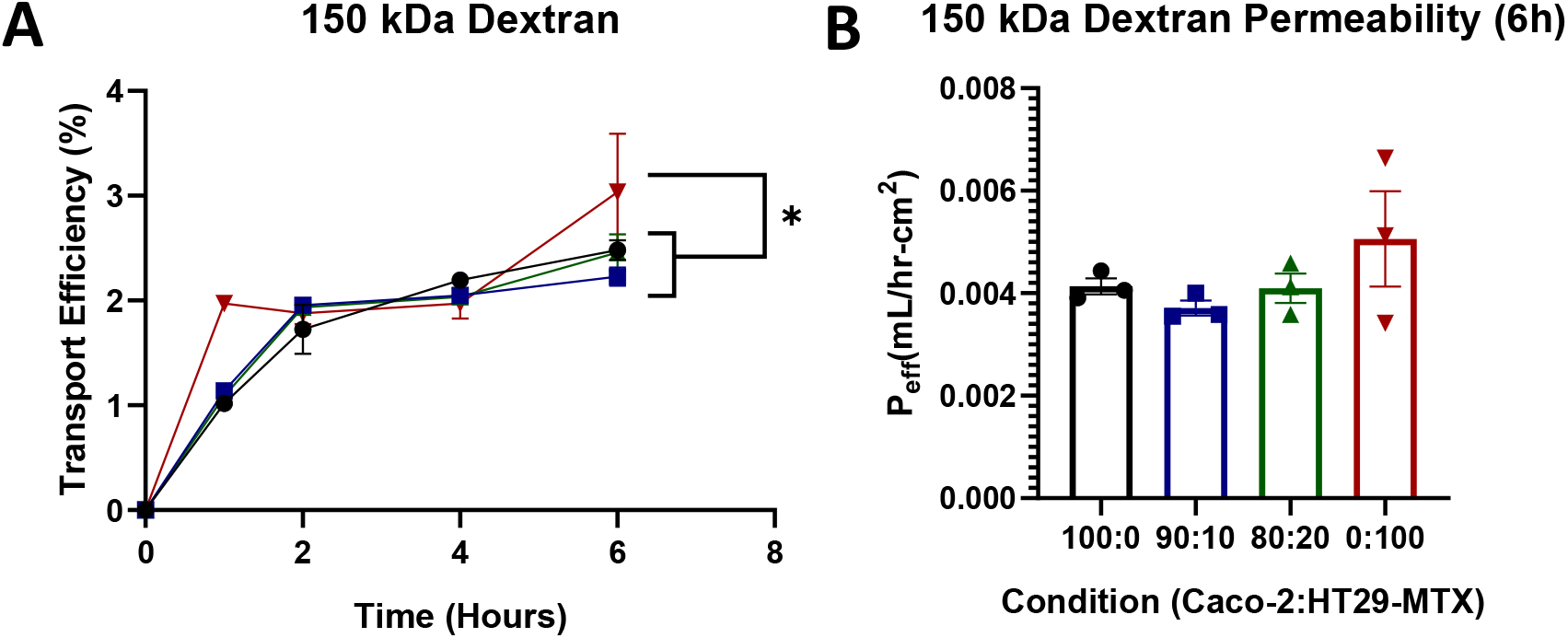
A) 150 kDa dextran transport efficiency across the transwell coculture over time after 3 weeks B) Effective permeability of cocultures at 6h to 150 kDa. *P < 0.05;) measured using 2-way ANOVA for transport efficiency over time and 1-way ANOVA for permeability.

