## Supplement for "Generating an *in vitro* gut model with physiologically relevant biophysical mucus properties"

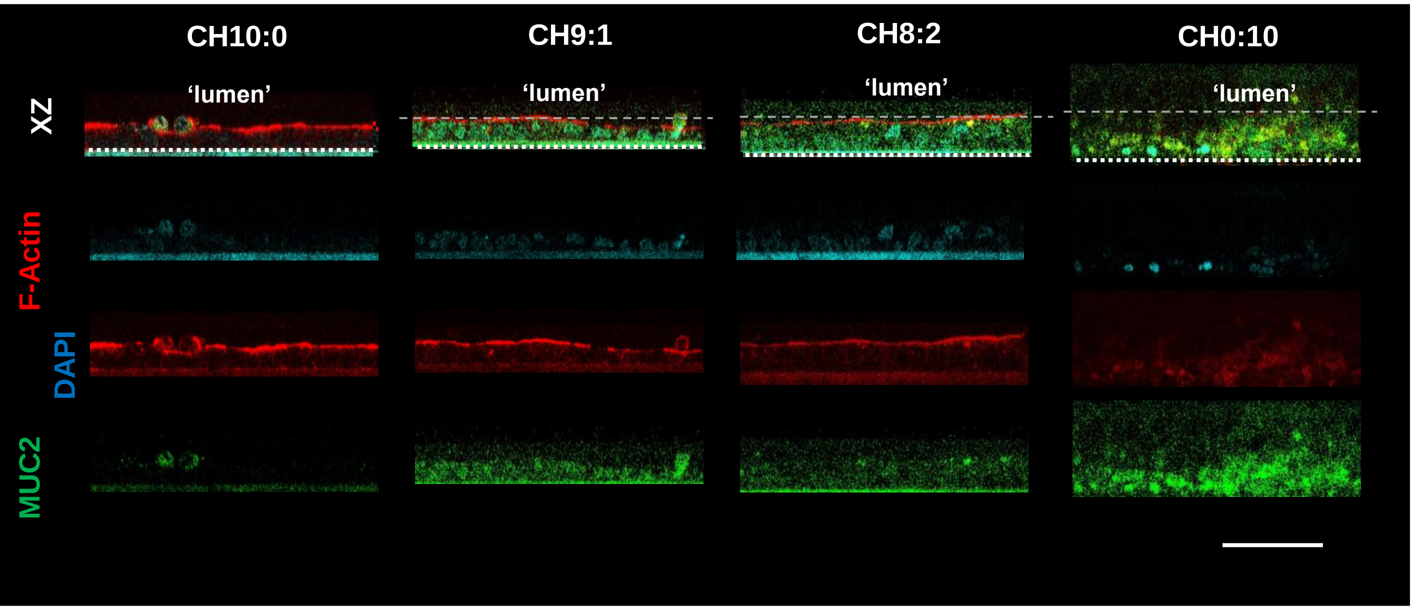


Supplemental Figure 1. A) Immunofluorescence images of different co-culture conditions visualizing cytoskeletal F-actin (Phalloidin, red), mucin (MUC2, green) and nuclei (DAPI, blue) Scale bar 200 µm.

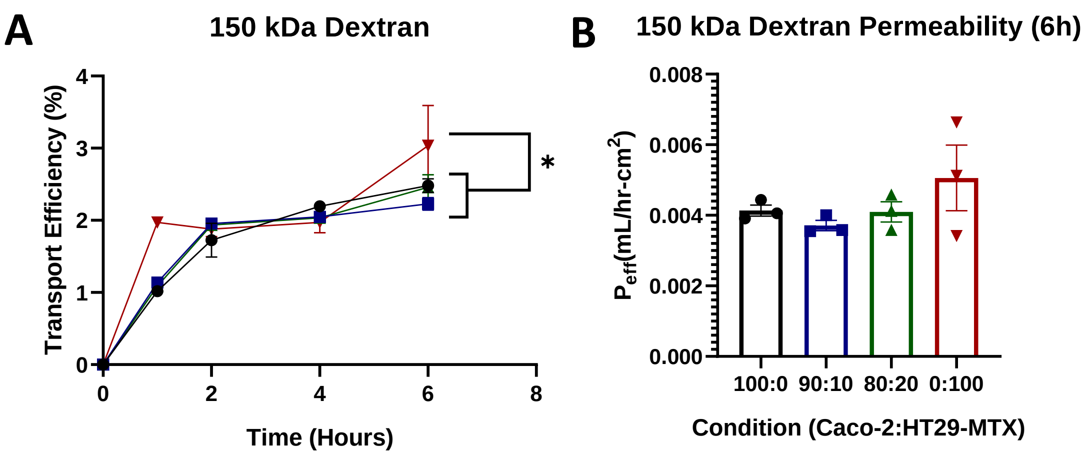


Supplemental Figure 2. A) 150 kDa dextran transport efficiency across the transwell coculture over time after 3 weeks B) Effective permeability of cocultures at 6h to 150 kDa. *$P < 0.05$;) measured using 2-way ANOVA for transport efficiency over time and 1-way ANOVA for permeability.
